# Development of a CRISPR/Cas9-induced gene editing system for *Pseudoalteromona fuliginea* and its applications in functional genomics

**DOI:** 10.1101/2025.05.30.657001

**Authors:** Zedong Duan, Ruyi Yang, Tingyi Lai, Wanning Jiang, Jin Zhang, Bo Chen, Li Liao

**Affiliations:** Key Laboratory for Polar Science, Ministry of Natural Resources, Polar Research Institute of China, Shanghai, China; Key Laboratory of Polar Ecosystem and Climate Change, Ministry of Education; Shanghai Key Laboratory of Polar Life and Environment Sciences; and School of Oceanography, Shanghai Jiao Tong University, Shanghai, 200030, China; School of Health Science and Engineering, University of Shanghai for Science and Technology, Shanghai, 200093, China

## Abstract

*Pseudoalteromonas* has been used as a model system to study cold adaptation and is of widespread interest in biotechnology and ecology. To explore its physiological responses to extreme cold, uncover functional genes, and clarify their ecological roles, efficient genetic tools are essential. However, existing genetic manipulation methods in *Pseudoalteromonas* rely on traditional homology-based recombination, which is inefficient and time-consuming. Consequently, improving editing efficiency is crucial for advancing both basic research and applied potential. Here, we introduced the CRISPR/Cas9 system into *Pseudoalteromonas* for the first time, and conducted an extensive investigation into the application of the Type II CRISPR/Cas9 system for gene editing in *Pseudoalteromonas fuliginea*, a representative species thriving in the frigid polar oceans. To validate the feasibility of the CRISPR/Cas system in *P. fuliginea*, multiple genes were selected as targets and confirmed the gene editing effects through phenotypic changes or gene expression. We have successfully achieved both gene knockouts and insertions in *P. fuliginea*, encompassing the deletion of genes such as *fliJ*, *indA*, and genes encoding Pf sRNAs, as well as the in *vivo* insertion of *3×flag* and the *gfp* gene. The average CRISPR/Cas9 gene editing efficiency in *P. fuliginea* exceeded 70% (range: 73.3%-95.8%), which is significantly higher than the traditional homology-based approach (less than 0.1%). In summary, we developed an efficient CRISPR/Cas9-based editing system in *P. fuliginea*, which can be utilized to accelerate the development of *Pseudoalteromonas* as a model system for addressing fundamental questions related to extreme environmental adaptation and to fulfill its potential biotechnological applications.

**IMPORTANCE:** *Pseudoalteromonas fuliginea* is a marine bacterium with great potential for ecological and biotechnological research, yet its genetic manipulation has long been a technical challenge. In this study, we developed a gene editing system based on CRISPR technology that enables efficient and precise genome modification in this organism. Using this system, we successfully deleted, inserted, and tagged multiple genes, including regulatory and non-coding elements, with high success rates. Notably, several of these genes are linked to key traits such as motility and stress response, which contribute to microbial adaptation in polar environments. This tool allows researchers to directly test gene function and study microbial adaptation in cold marine environments. The ability to perform reliable genetic edits in *Pseudoalteromonas fuliginea* opens new possibilities for its use as a model organism and will support future advances in microbial ecology, environmental microbiology, and marine biotechnology.

## Introduction

*Pseudoalteromonas* is a genus of marine bacteria within the class Gammaproteobacteria, known for its ecological relevance and biotechnological potential. Members of this genus account for approximately 0.5- 6.0% of the global marine planktonic bacterial communities and are particularly abundant in polar regions such as the Arctic and Southern Oceans (1). Strains of *Pseudoalteromonas* have been isolated from diverse marine environments, including seawater, deep-sea sediments (2), sea ice (3), and a variety of eukaryotic hosts such as macroalgae (4), sponges (5) and marine animals (6). These features make *Pseudoalteromonas* a valuable model system for studying microbial adaptation to extreme environments (7). The ecological significance of *Pseudoalteromonas* is further highlighted by its interactions with marine algae and animals. Some strains can promote algal growth, while others exhibit inhibitory effects (8). Meanwhile, *Pseudoalteromonas* is a prolific producer of bioactive compounds, including extracellular polysaccharides, pigments, and antimicrobial substances (9–12). These bacteria are also a rich source of cold-active enzymes, making them highly attractive for applications in environmental microbiology, biotechnology, and biomedicine.

To advance both fundamental and applied research on *Pseudoalteromonas*, efficient genetic manipulation tools are essential. Unlike model organisms such as *Escherichia coli*, genetic engineering in *Pseudoalteromonas* remains underdeveloped. While several plasmids have been introduced, their application is limited in scope and efficiency. For instance, Zhao et al. discovered a native plasmid, pSM429, in *Pseudoalteromonas sp*. BSi20429, which is 3874bp in size and has a GC content of 28% (13). By integrating mobile elements, they constructed the shuttle plasmid pWD, enabling conjugative transfer between *E. coli* and *Pseudoalteromonas*. This shuttle plasmid was successfully used to express the erythromycin resistance gene in the host. Similarly, the endogenous plasmid pMtBL from *P. haloplanktis* TAC 125 was modified to create a shuttle vector for heterologous protein expression (14). Wang et al. employed pK18*mobsacB*-Ery and pK18*mobsacB*-Cm plasmids to achieve in-frame deletion of genes in several *Pseudoalteromonas* species (15). Other plasmids, such as pVS (16) and pMT (17) have also been applied for gene editing in this genus.

However, these tools largely rely on traditional double-crossover homologous recombination, which is in general inefficient, time-consuming, and labor-intensive. As a result, genetic manipulation remains a major bottleneck limiting the functional analysis of *Pseudoalteromonas*. There is a clear and urgent need for a more versatile, rapid, and efficient system that can support sequential gene edits and broader genetic modifications. Such advances would significantly accelerate research on gene function and facilitate the broader application of *Pseudoalteromonas* in microbial ecology and biotechnology.

The CRISPR/Cas system is a widely used gene editing tool, valued for its efficiency, ease of use, and broad applicability. It has been successfully implemented in a variety of organisms, including *E. coli*, *Streptomyces*, *Saccharomyces cerevisiae*, higher plants, and human cell lines (18–24). Despite its widespread use, CRISPR/Cas has not previously been introduced in *Pseudoalteromonas*. Several factors influence the feasibility of CRISPR/Cas implementation in a new species, including the availability of efficient DNA delivery methods, expression of Cas proteins and guide RNAs, compatibility of Cas variants, and the host’s DNA repair mechanisms—particularly homologous recombination and non-homologous end joining (NHEJ). In this study, we have developed an efficient CRISPR/Cas9 gene editing system tailored specifically for *P. fuliginea* to enhance the effectiveness of existing genetic manipulation. Building on the plasmid backbone pK18mobsacB-Ery, we constructed a series of CRISPR/Cas9-based vectors, incorporating optimized Cas9 expression elements and suitable promoters compatible with *P. fuliginea*. These vectors enabled us to achieve targeted gene knockouts, insertions, and knockdowns with high precision. To evaluate the system’s performance, we applied it to multiple genetic targets and verified editing outcomes through phenotypic analysis and protein expression. The results demonstrated that our CRISPR/Cas9 platform consistently achieved editing efficiencies above 70%, representing a substantial improvement compared to traditional double-crossover homologous recombination approaches.

## Results and Discussion

### Construction of CRISPR/Cas9 plasmid for *P. fuliginea*

To develop a functional CRISPR/Cas9 editing system for *P. fuliginea*, we constructed a recombinant plasmid based on pK18mobsacB-Ery, a shuttle vector that has been successfully used for gene knockout in *Pseudoalteromonas* via double-crossover homologous recombination, and is capable of conjugative transfer between *E. coli* and *Pseudoalteromonas* (15). The CRISPR/Cas9 cassette was assembled by inserting the Cas9 coding sequence and a single guide RNA (sgRNA) expression module into the pK18mobsacB-Ery backbone. For efficient gene expression in *P. fuliginea*, we screened suitable promotors to drive both Cas9 and sgRNA expression. As a result, the D4-12 sequence of the Pf1 sRNA promoter was selected, as it had previously been demonstrated to be an active regulatory element in *P. fuliginea* BSW20308 (25). Its native origin increases the likelihood of strong transcriptional activity in this host. To ensure proper transcription termination, the rrnB T1 sequence was employed as a rho-independent terminator downstream of both expression units. This element has been widely used in prokaryotic expression systems due to its robust and reliable termination efficiency, which prevents transcriptional read-through (26). The Cas9 protein used in this system was derived from *Streptococcus pyogenes* MGAS5005 (27) and was codon-optimized to match the codon usage preferences of *Pseudoalteromonas*, thereby improving translation efficiency in the heterologous host. The complete expression cassette, which includes the D4-12 promoter, codon-optimized Cas9, sgRNA module, and *rrnB* T1 terminator, was cloned into the pK18mobsacB-Ery plasmid to yield the final CRISPR/Cas9 construct (Figure 1A). To validate the CRISPR/Cas9 system in *P. fuliginea*, we performed gene editing on the *P. fuliginea* BSW20308 strain, as detailed in the subsequent sections.

**Figure 1.**
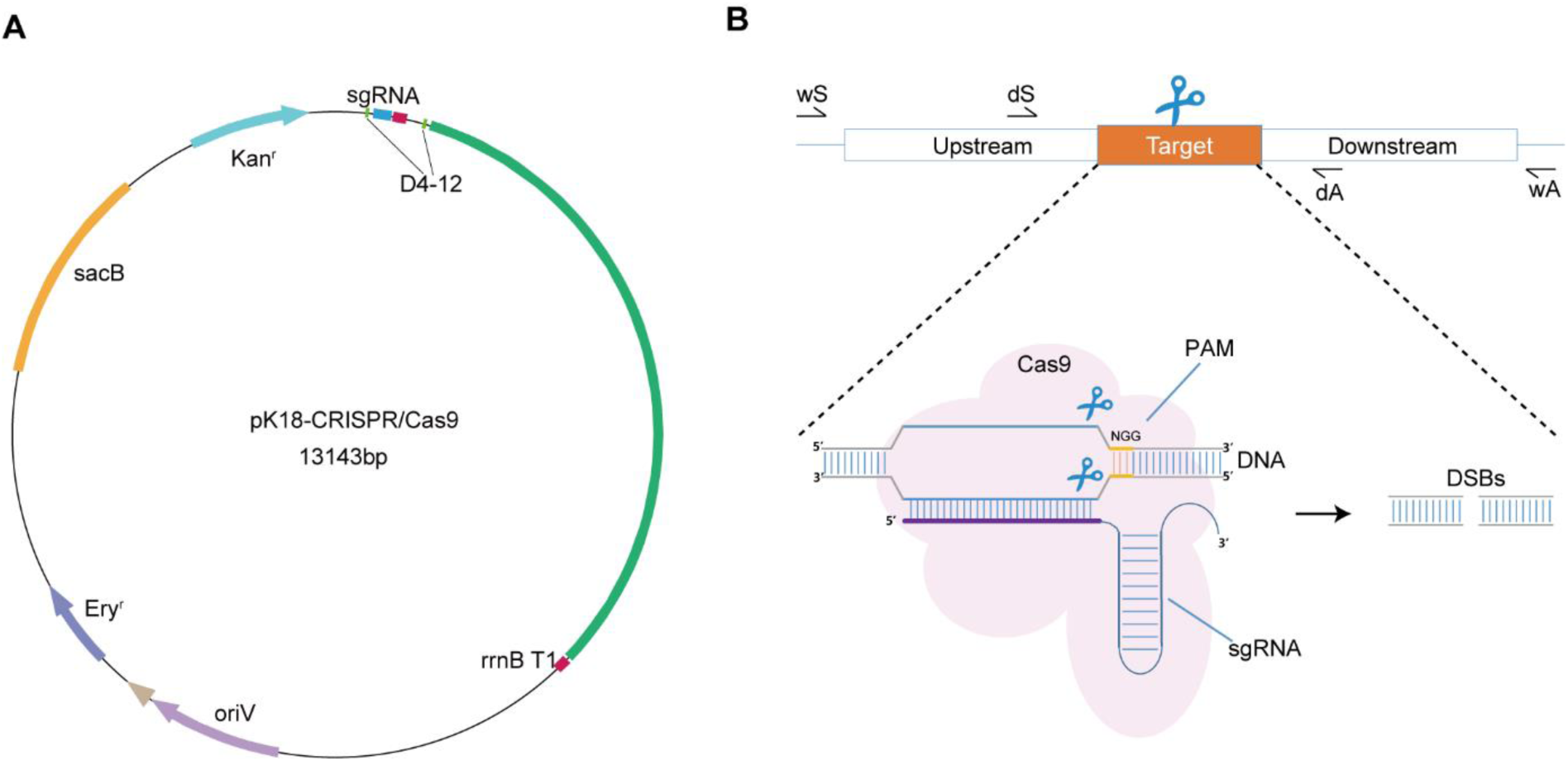
The construction of pK18Ery-CRISPR/Cas9 for gene editing in *Pseudoalteromona*. (**A**) Map of the pK18Ery-CRISPR/Cas9 mobilizable shuttle vectors. (**B**) Guided by the gRNA, the Cas9 protein induces a double-strand break in the DNA at the PAM (Protospacer Adjacent Motif) site.

### Deletion of motility gene *fliJ* in *P. fuliginea* through CRIPSR/Cas9 system

The *fliJ*, integral to the bacterial flagellar structure and pivotal for cell motility and chemotaxis, is present within the genome of *P. fuliginea* BSW20308 (28). Consequently, we opted to eliminate this gene to assess the efficacy of the CRISPR/Cas9 system in *P. fuliginea*. A CRISPR/Cas9 suicide plasmid, pK18Ery- fliJ-CRISPR/Cas9, was constructed based on the pK18*mobsacB*-Ery backbone. This plasmid carried the Cas9 gene, a specific sgRNA targeting *fliJ*, and two homologous arms (1,000 bp each) flanking the *fliJ* locus. Conjugative transfer of the plasmid into *P. fuliginea* BSW20308 was performed, and transconjugants were selected on 2216E agar containing erythromycin. Following successful transfer, the Cas9 protein was expressed under the control of the D4-12 promoter. The Cas9-sgRNA complex induced a double-strand break at the target site adjacent to the PAM sequence (Figure 1B), which was repaired via homologous recombination using the provided flanking sequences, resulting in the deletion of the target gene.

To confirm the gene knockout, we performed PCR using a primer pair (*fliJ*-wS/ *fliJ*-dA) spanning the upstream homologous region and the 3’ end of the *fliJ* locus. The deletion of the *fliJ* gene (447 bp) was confirmed by gel electrophoresis (Figure 2A) and further validated through DNA sequencing. Out of 17 randomly selected erythromycin-resistant clones, 15 exhibited successful gene deletions, corresponding to an editing efficiency of 88% (Figure S1, Supplementary Data 1). In contrast, when attempting to delete *fliJ* using the traditional double-crossover homologous recombination method, which utilized the same plasmid but lacked the CRISPR/Cas cassette, none of the 100 screened colonies yielded a successful mutant. This outcome underscores the markedly superior efficiency of the CRISPR/Cas9 approach. To further validate the successful genetic manipulation via phenotypic analysis, cell motility assays were performed on 0.3% agar plates. The Δ*fliJ* mutant exhibited a significant reduction in motility compared to the wild-type strain, aligning with the anticipated role of *fliJ* in flagellar assembly (Figure 2B). This impaired motility may reduce the strain’s ability to respond to environmental stimuli or colonize new niches, highlighting the role of fliJ in microbial adaptation to physically challenging or nutrient-scarce environments such as polar oceans.

**Figure 2.**
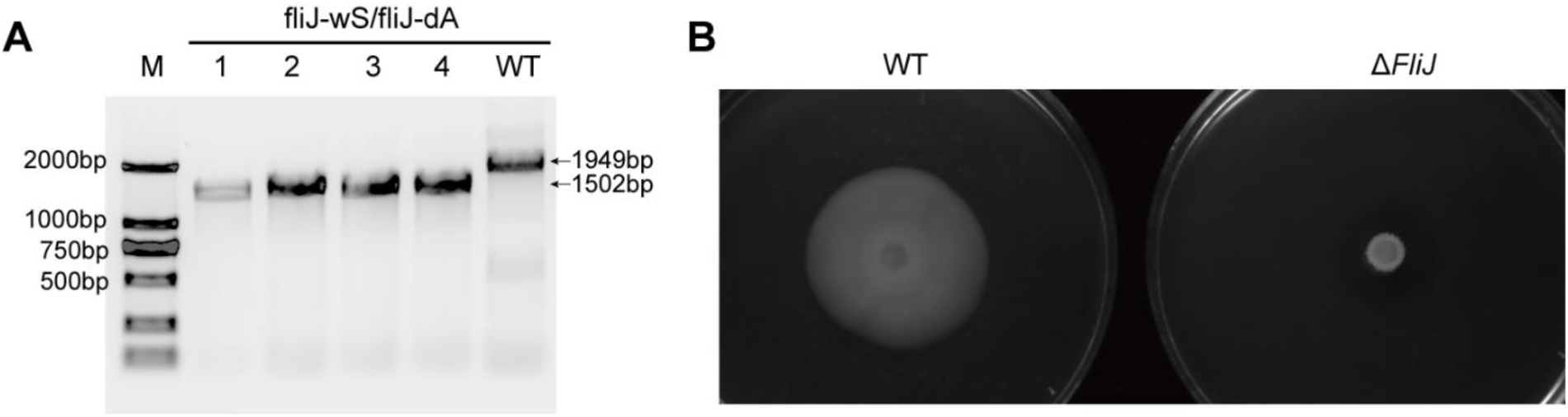
Deletion of motility gene *fliJ* in *P. fuliginea* BSW20308 through CRIPSR/Cas9 system. (**A**) PCR confirmation of complete deletion of the *fliJ* gene using the *fliJ*-wS and *fliJ*-dA primers. 1-4, four independent clones after the successful conjugation of the pK18Ery-*fliJ*-CRISPR/Cas9 suicide plasmid. WT indicates BSW20308 wild-type strain. (**B**) Swimming ability of the WT and Δ*FliJ* strains indicated by spreading area on 0.3% agar plates.

### Deletion of indigoidine biosynthetic gene *indA* in *P. fuliginea* through CRISPR/Cas9

*P. fuliginea* BSW20308 is capable of producing a characteristic dark blue pigment when grown on 2216E medium. Previous research identified a potential indigoidine biosynthetic gene cluster in this strain, which includes the *indA* gene as a candidate gene likely responsible for pigment production (28). To investigate the function of *indA* and further validate the CRISPR/Cas9 system, we constructed a deletion mutant targeting this gene. Homologous fragments of 1,000 bp upstream and downstream of *indA* were inserted into the pK18Ery-indA-CRISPR/Cas9 plasmid. The resulting construct was introduced into *P. fuliginea* BSW20308 via conjugation, and integration was confirmed by PCR. Subsequently, counter-selectable 2216E media containing 10% sucrose was utilized to product the Δ*indA* mutant. The PCR product of Δ*indA* was 310 bp, while the corresponding wild-type PCR product was 1255 bp (Figure 3A). DNA sequencing further confirmed the deletion of the 945 bp of *indA* coding region. To assess the phenotypic effects of this deletion, we compared pigment production between wild-type and mutant strains. The Δ*indA* mutant lost the ability to synthesize the characteristic blue pigment, confirming the functional role of *indA* in indigoidine biosynthesis (Figure 3B). Moreover, functional complementation was performed by introducing a plasmid carrying the intact *indA* gene (pK18-indA) into the Δ*indA* mutant. The complemented strain regained the wild-type phenotype and restored pigment production, providing further evidence that the loss of pigment was directly attributable to *indA* deletion. In total, 24 erythromycin-resistant colonies were screened, of which 19 exhibited successful *indA* gene deletions, corresponding to a gene editing efficiency of approximately 79% (Figure S2).

**Figure 3.**
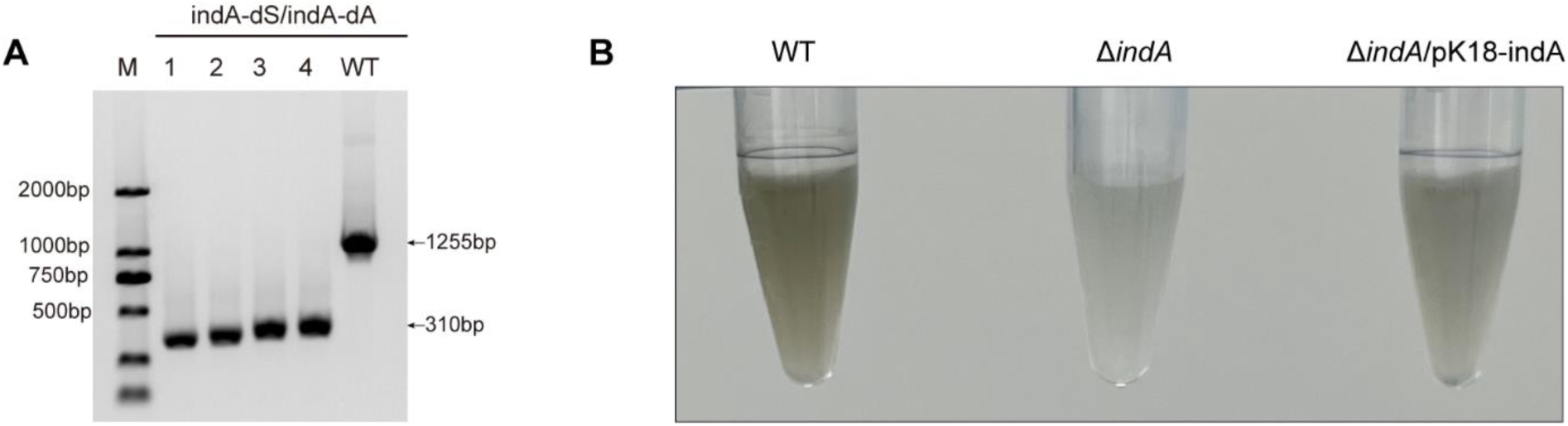
**Deletion of indigoidine biosynthetic gene *indA* in *P. fuliginea* through CRISPR/Cas9**. (**A**) PCR confirmation of *indA* deletion using the *indA*-dS and *indA*-dA primers. 1-4, four independent clones after the successful conjugation of the pK18Ery-*indA*-CRISPR/Cas9 suicide plasmid. WT indicates BSW20308 wild-type strain. (**B**) The pigment production of wild-type *P. fuliginea* BSW20308, Δ*indA* and Δ*indA*/pK18-*indA*.

### Stepwise cumulative CRISPR/Cas9-mediated gene knockout of three Pf sRNA loci in *P. fuliginea*

In a previous study, an attempt was made to delete the small RNA gene *Pf1* through double-crossover homologous recombination. However, only one mutant was recovered from nearly a thousand colonies screened, resulting in an extremely low efficiency of approximately 0.1% (25). To address this limitation, we constructed CRISPR/Cas9-based plasmids targeting three small RNAs (*Pf1*, *Pf2*, and *Pf3*), designated as pK18Ery-Pf1-CRISPR/Cas9, pK18Ery-Pf2-CRISPR/Cas9, and pK18Ery-Pf3-CRISPR/Cas9, respectively. These plasmids were mobilized into *P. fuliginea* BSW20308 in a sequential manner. The pK18Ery-Pf1- CRISPR/Cas9 plasmid was first introduced into BSW20308 via conjugation. The successful deletion of *Pf1* was confirmed by PCR and DNA sequencing (Figure S3). Using the Δ*Pf1* mutant as a starting strain, we next knocked out *Pf2* to generate the Δ*Pf12* double mutant, followed by deletion of *Pf3* to obtain the Δ*Pf123* triple mutant. The editing efficiencies achieved for *Pf1*, *Pf2*, and *Pf3* using the CRISPR/Cas9 system were 92%, 86.7%, and 73.3%, respectively (Figure S3, Supplementary Data 1). These values represent a substantial improvement over the homologous recombination method and highlight the effectiveness of our CRISPR/Cas9 platform in targeting non-coding RNA genes in *P. fuliginea*.

### *csrA* knock-down in *P. fuliginea* through CRIPSPR/Cas9 system

The CsrA protein is a global post-transcriptional regulator involved in diverse cellular processes through RNA binding, and plays a central role in the carbon storage regulator (Csr) system. To investigate its function in *P. fuliginea*, we initially attempted to knock out the *csrA* gene. However, repeated attempts failed to produce viable knockout mutants, suggesting that *csrA* may be essential for cell survival under standard growth conditions (29, 30). To circumvent this limitation, we constructed a modified plasmid, pK18Ery- *csrA_1-50_*-CRIPSPR/Cas9, which included a donor fragment encoding the first 50 amino acids of CsrA. This design aimed to generate a partial deletion or functional disruption of *csrA* while preserving minimal functionality to maintain cell viability. The plasmid was mobilized into *P. fuliginea* via conjugation, and positive transconjugants were identified using two sets of plasmid-specific primers (Ery-F/Ery-R and SacB-F/SacB-R). Sequencing of the *csrA* locus in edited clones revealed a 39 bp deletion at the 3’ end of the gene (Figure 4), demonstrating successful knockdown. Among the screened colonies, the CRISPR/Cas9-mediated knockdown efficiency for *csrA* was approximately 25%. This relatively low efficiency is likely due to the essential role of *csrA* as a global post-transcriptional regulator. Disrupting this gene may impair key cellular functions, making complete or partial edits detrimental to cell survival under standard conditions.

**Figure 4.**
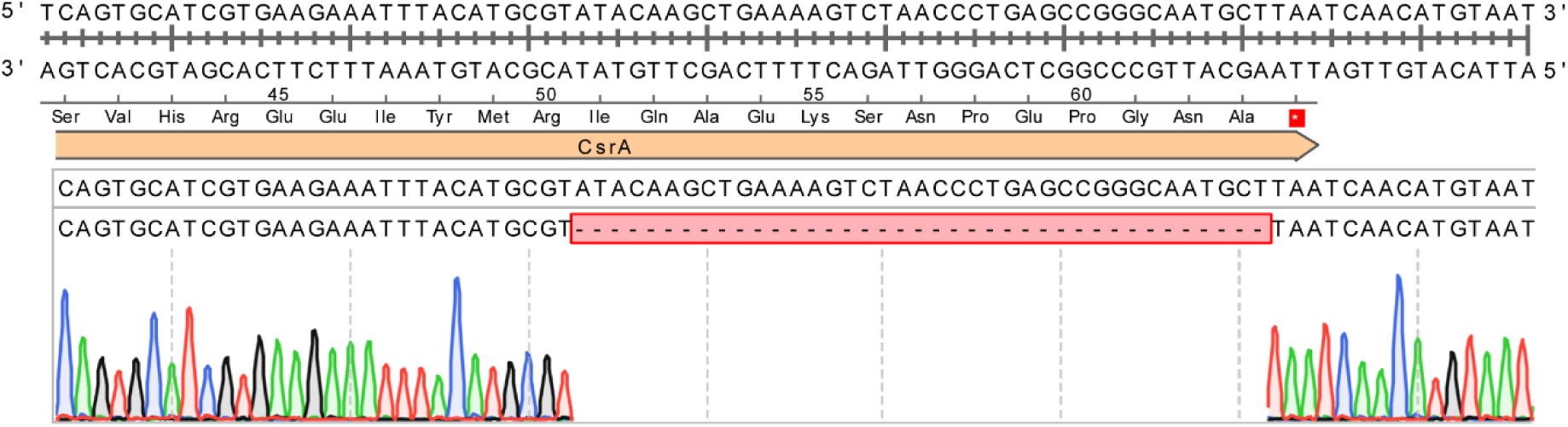
***csrA* knock-down in *P. fuliginea* through CRIPSPR/Cas9 system**. Sanger sequencing of purified PCR products confirmed knock down of *csrA* gene.

### CRISPR/Cas9-induced gene knock-in for *P. fuliginea*

To evaluate the gene knock-in capability of the CRISPR/Cas9 system in *P. fuliginea* BSW20308, we engineered the insertion of a 3×FLAG epitope tag at the 3’ end of the *csrA* gene. A recombinant plasmid, pK18Ery-*csrA*::*3×flag*-CRISPR/Cas9, was constructed by assembling the *csrA* sequence, the 3×FLAG tag, and homologous arms flanking the *csrA* locus, along with the guide RNA cassette and Cas9 gene, into the pK18Ery backbone. This plasmid was then introduced into *P. fuliginea* via conjugation. Entry of the plasmid was confirmed using plasmid-specific primer pairs (Ery-F/Ery-R and SacB-F/SacB-R). Successful knock-in of the 3×FLAG sequence was confirmed by PCR using primers FLAG-KI-dS and FLAG-KI-wA. The resulting 1,141 bp amplicon was detected in edited mutants but not in the wild-type strain (Figure 5A). DNA sequencing confirmed precise insertion of the 3×FLAG tag at the 3’ terminus of the *csrA* gene. Protein-level expression of the CsrA-3×FLAG fusion product was validated via western blot using a FLAG-specific antibody. The results showed detectable expression of the CsrA-3×FLAG fusion protein in the knock-in strains (Figure 5B), further validating the insertion.

**Figure 5.**
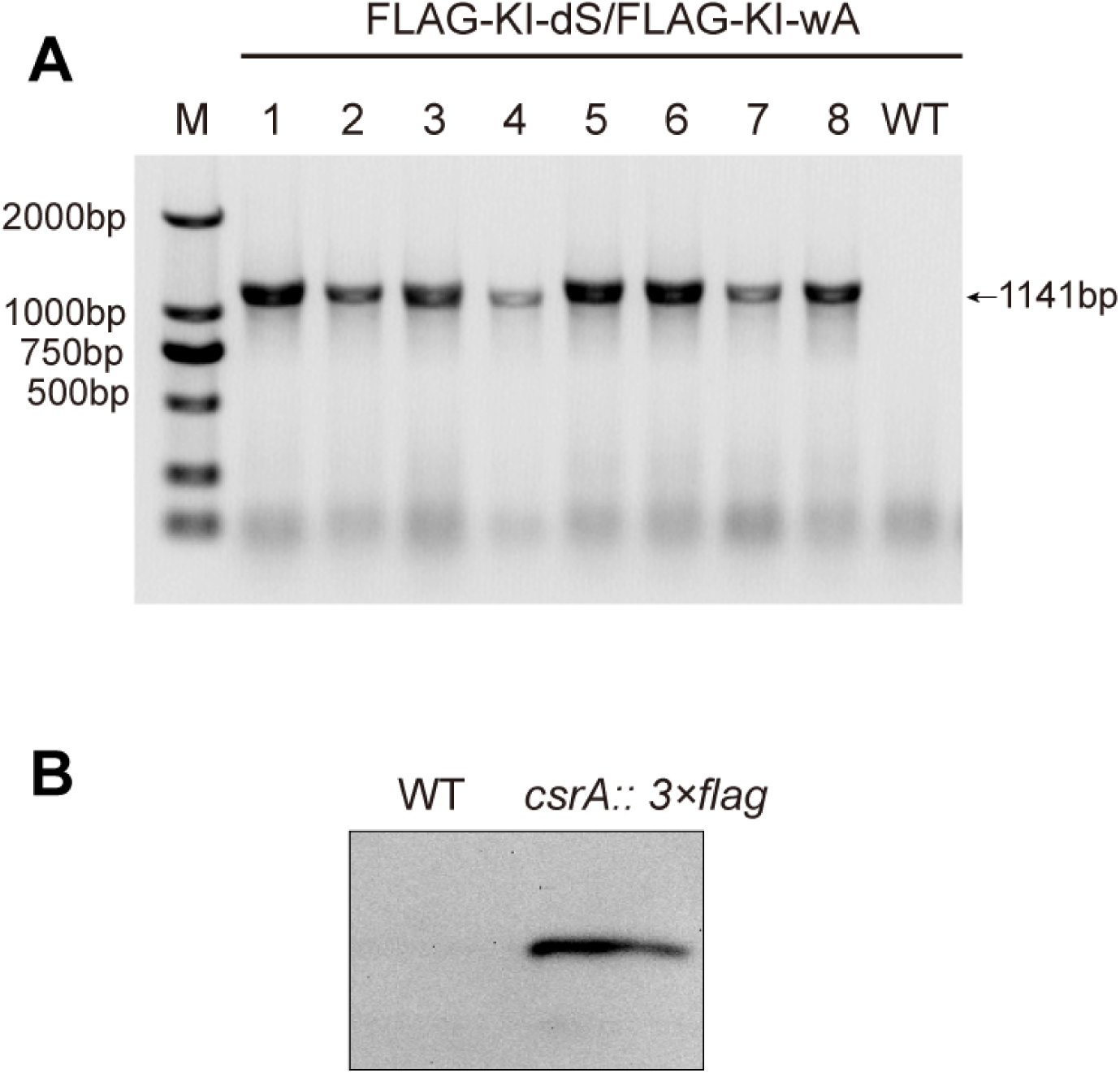
CRISPR/Cas9-induced gene knock-in for *P. fuliginea*. (**A**) PCR confirmation of *3×flag* insertion using the *fliJ*-wS and *fliJ*-dA primers. 1-8, eight independent clones after the successful conjugation of the pK18Ery-*csrA*::*3×flag*-CRISPR/Cas9 suicide plasmid. WT indicates BSW20308 wild-type strain. (**B**) Expression of 3*×*FLAG tag was detected in cell lysates of the indicated *P. fuliginea* strains using western blot. WT, wild type strain.

Editing efficiency was assessed by screening 48 erythromycin-resistant clones through PCR. Of these, 46 showed successful amplification of the 3×FLAG tag. To eliminate the possibility of false negatives, the two non-amplified clones were retested in triplicate, all of which yielded no detectable bands. This corresponds to a knock-in efficiency of 95.8% (Figure S4), demonstrating the high precision and reliability of CRISPR/Cas9-mediated gene tagging in *P. fuliginea*.

### Targeted insertion of a fluorescent reporter at the *tse2* locus using CRISPR/Cas9 in *P. fuliginea*

To further demonstrate the versatility of the CRISPR/Cas9 system in *P. fuliginea*, we performed targeted gene tagging with a fluorescent reporter. A gene encoding a protein with a conserved Tse2 ADP-ribosyltransferase (ADPRT) toxin domain was selected as the insertion site for a green fluorescent protein (GFP) reporter(31), providing a useful model for downstream applications. A donor construct was designed by appending the *gfp* gene to the 3’ end of the *tse2* coding sequence within the pK18Ery-*tse2::gfp*- CRISPR/Cas9 plasmid. This recombinant plasmid, carrying homologous arms flanking the *tse2* gene, Cas9, and the sgRNA cassette, was mobilized into *P. fuliginea* via conjugation. Successful integration of the *gfp* sequence at the 3’ end of the target gene was verified by PCR and confirmed through DNA sequencing (Figure 6A). Fluorescence microscopy revealed detectable, albeit weak, GFP signal in the modified strain. This observation was further supported by fluorescence quantification, confirming expression of the Tse2-GFP fusion protein (Figure 6B). Among 24 erythromycin-resistant clones screened, 20 showed successful insertion of the *gfp* gene, corresponding to a gene editing efficiency of 83.3% (Figure S5). These results validate the feasibility of using CRISPR/Cas9 for targeted fluorescent tagging in *P. fuliginea*, offering a valuable tool for real-time protein tracking and functional analysis in this organism.

**Figure 6.**
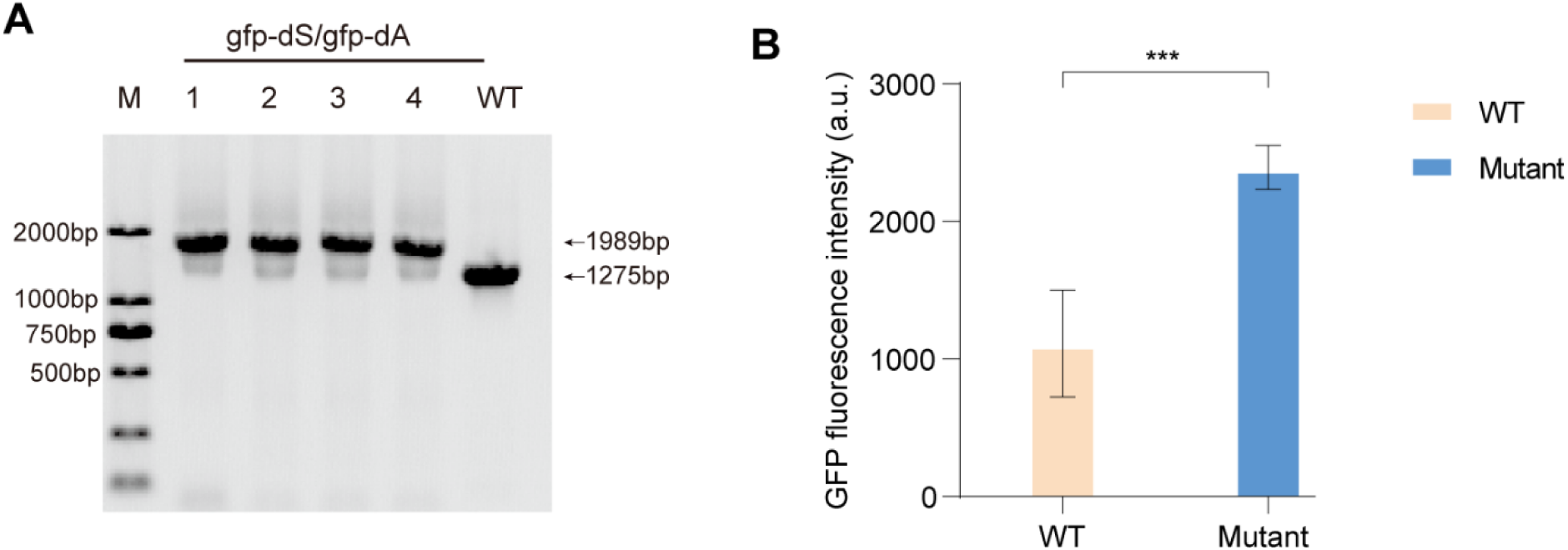
Insertion of *gfp* gene in *P. fuliginea* genome through CRISPR/Cas9. (**A**) PCR confirmation of *gfp* insertion using the *gfp*-dS and *gfp*-dA primers. 1-4, four independent clones after the successful conjugation of the pK18Ery-*tse2::gfp*-CRISPR/Cas9 suicide plasmid. WT indicates BSW20308 wild-type strain. (**B**) GFP fluorescence measurements to assess Tse2-GFP fusion protein expression in *P. fuliginea*. Plotted is the mean ± s.e.m (\*\*\**P* < 0.001 using Student’s t-test).

## Conclusions

This study presents the first successful establishment of a CRISPR/Cas9-based genome editing system for *P. fuliginea*. By constructing a series of optimized editing plasmids, we demonstrated that this system enables efficient gene knockouts, knockdowns, and knock-ins with high reliability and versatility. Across multiple genomic targets—including structural genes (*fliJ*), biosynthetic genes (*indA*), regulatory elements (*csrA*), small RNAs (*Pf1–Pf3*), and tagged reporter constructs (*csrA::3×flag* and *tse2::gfp*)—editing efficiencies consistently exceeded 70% except for the relatively low knock-down efficiency of the *csrA* gene. Compared to traditional double-crossover recombination, which showed low success in similar editing attempts, the CRISPR/Cas9 system provided a dramatic improvement in both efficiency and workflow simplicity. Overall, this CRISPR/Cas9 platform provides a powerful and flexible genetic toolkit for *P. fuliginea*, greatly enhancing its potential as a model organism for studying cold adaptation, regulatory networks, and host-microbe interactions. It also lays a technical foundation for future applications in marine biotechnology and synthetic biology involving polar marine bacteria.

## Methods and Materials

### Strains, plasmids, and growth conditions

The strains and plasmids used in this study are listed in Supplementary Data 1. *E. coli* WM3064 strains were grown in LB medium with 0.3mM DAP (diaminopimelic acid) at 37℃ (10g tryptone, 5g yeast extract, and 10g NaCl dissolved in 1000 ml of deionized water), supplemented with kanamycin (50 mg/L) or gentamicin (10 mg/L). The *P. fuliginea* BSW20308 and its mutation strains were grown in 2216E medium. The plasmids used in this study are listed in Supplementary Data 1. The antibiotics were added at the following concentrations: 50 μg/ml for kanamycin (Kan); 25 μg/ml for erythromycin (Ery).

### Conjugation assays

The plasmids were transferred into *P. fuliginea* through conjugation, following the established protocols described in previous reports (15, 32). In brief, *E. coli* WM3064 (donor) and recipient strains were cultured to an OD_600_ of 0.6-1.0. The donor and the recipient strains were collected by centrifugation at 12,000 rpm, and washed twice with MLB medium (Modified LB medium containing 10g tryptone, 5g yeast extract, and 10g NaCl dissolved in 500 ml deionized water and 500 ml sterile seawater). The combined strain mixture was resuspended in 100 µl of MLB medium and then dropped onto MLB plates supplemented with diaminopimelic acid (DAP). The plates were incubated at 25°C for 24 hours. The bacterial lawn was scraped off, washed multiple times, and then spread onto erythromycin-containing 2216E plates to select the positive transconjugants.

### Construction of the mutants

The design of gRNA was carried out on the website (https://zlab.bio/guide-design-resources) in this study. The appropriate PAM type and host genome were selected through the "Cas-Designer" tool within the website. The target gene sequence that needed to be edited was then input into the "Target Sequence" field according to the specified format. After submission, the results were filtered based on requirements. The website featured a built-in scoring function, and typically, the gRNA with a higher score was selected for further use. The donor fragments, gRNA cassette, and Cas9 gene were ligated into the plasmid pK18*mobsacB*-Ery digested with *BamH* I and *Hind* III using the ClonExpress II one-step cloning kit (Vazyme Co., Ltd., China). Suicide CRISPR/Cas plasmid was transformed into *E. coli* WM3064, and then mobilized into *P. fuliginea* via conjugation. The positive transconjugants were screened by 2216E plates containing erythromycin (25 μg/ml), and confirmed by PCR followed by DNA sequencing. The mutant strains were spread onto 2216E solid medium containing 15% sucrose and incubated statically at 25°C until single colonies emerged. This process was repeated for 2-3 generations to ensure stability of the mutation. Subsequently, single colonies were picked and inoculated into 2216E liquid medium containing 15% sucrose for activation. To verify the loss of the plasmid, PCR was performed using two sets of primers specific to the plasmid fragments as amplification primers. The absence of PCR products corresponding to the plasmid-specific fragments would indicate successful loss of the plasmid.

### Cell motility assays

Strains were cultured to the exponential growth phase (OD_600_ of 0.4), centrifuged and resuspended in fresh 2216E medium for motility and biofilm assays. Cell motility was measured using semisolid agar plates in triplicate. For each strain, 5 μl of re-suspended cells were placed in the center of a 2216E plate containing 0.3% agar and incubated at 32°C for over 24 h.

### Western Blot

For western blot analysis, samples were transferred from the SDS-polyacrylamide (PAA) gels onto the polyvinylidene difluoride (PVDF) membranes be electroblotting. The PVDF membranes were blocked for 2 h with 50 ml of 5% (w/v) milk powder that dissolved in PBST (PBS with 0.5% Tween-20), and then incubated overnight at 4°C with primary antibody (Anti-FLAG Tag monoclonal antibody, Sangon Biotech, #D191041, China). Membranes were then washed with PBST solution for three times, followed by incubation with secondary antibody (HRP-conjugated Goat anti-mouse IgM, Sangon Biotech, #D110103, China) for 30 min. After washing with PBST for three times, the blot was developed using High sensitive Plus ECL luminescence reagent (Sangon Biotech, #C520045, China) and visualized using ChemiDoc MP Imaging System (Bio-Rad, USA). The strains with native CsrA were used as the negative controls.

### GFP expression in *P. fuliginea* and measurement of GFP fluorescence

The WT and *tse2::gfp* mutant were grown in 2216E medium, cells were collected by removing the medium, washed three times, and resuspended in 200 µl of sterile distilled water. The WT strain was used as the negative control. Five biological replicates were performed for both the control and experiment groups. The GFP fluorescence intensity was measured using a microplate reader (excitation, 485nm; emission, 538nm) and divided by the corresponding OD_600_ value to determine the unit fluorescence intensity.

## Acknowledgements

The authors thank Professor Xiulan Chen at Shandong University for kindly providing the pK18*mobsacB*-Ery plasmids. The project was supported by the National Natural Science Foundation of China (Grant No. 42476264 and 41976224) and the National Key Research and Development Program of China (Grant No. 2022YFC2807501).

